## Supplementary material for "Optimization of AAV tools to target Müller glial cells for retinal gene therapy": Figure S1-S5

**Supplemental figures**

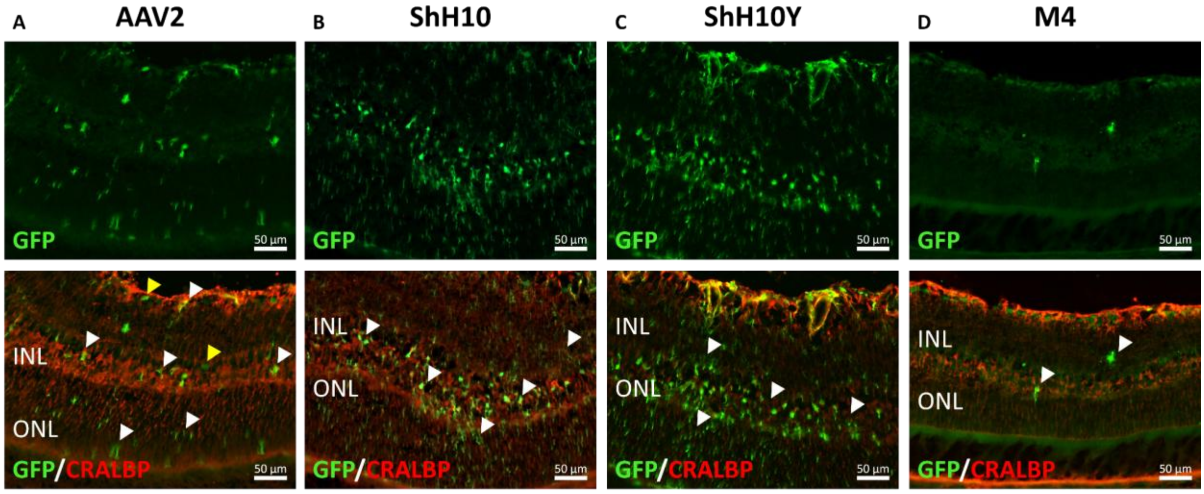


**Figure S1: MG marker analysis in rat retinae treated with different AAV capsids**. Representative images showing the expression pattern of GFP and the MG marker CRALBP in rat retinae treated with the AAV serotypes A) AAV2, B) ShH10, C) ShH10Y and D) M4 in combination with the *GFAP* promoter. Scale bar = 50 µm.

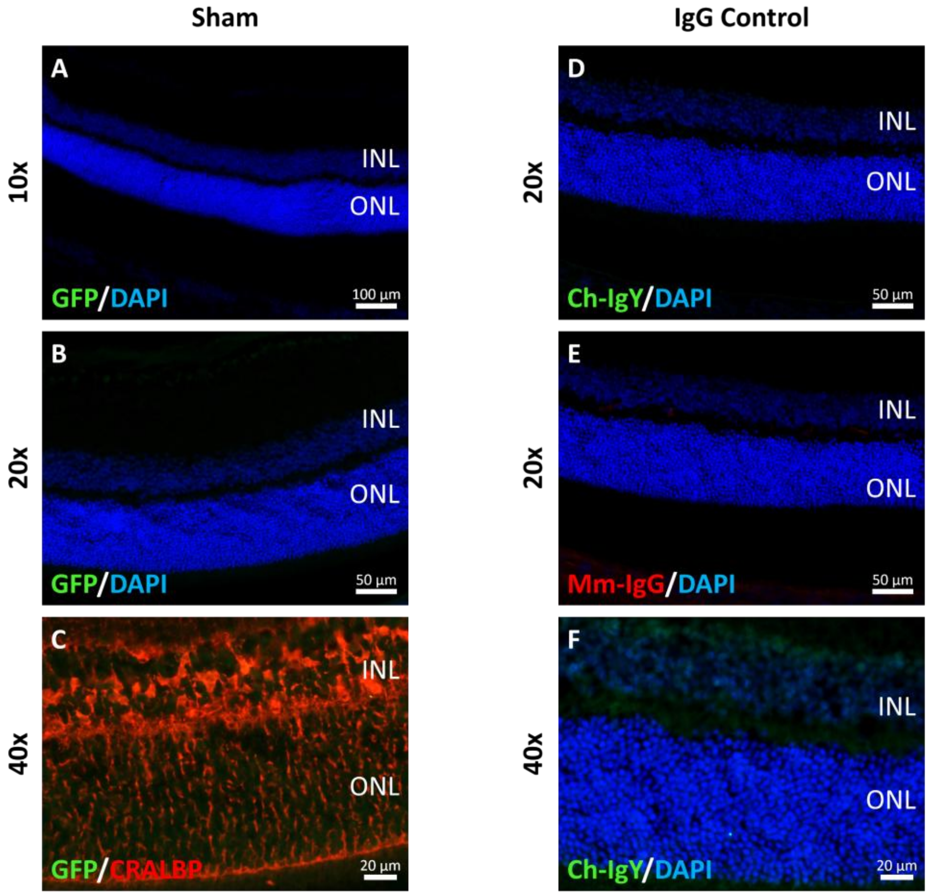


**Figure S2: Controls for the *in vivo* AAV transduction analysis.** A-C) Representative images of immunohistochemistry analysis to detect GFP and CRALBP expression in rat retinae injected with saline solution (sham control). Scale bar = 100 µm (A); 50 µm (B); 20 µm (C). D-F) Representative images showing clean background in the IgG control staining for Chicken (Ch-IgY) and mouse (Mm-IgG) antibodies. Scale bar = 50 µm (D-E); 20 µm (F).

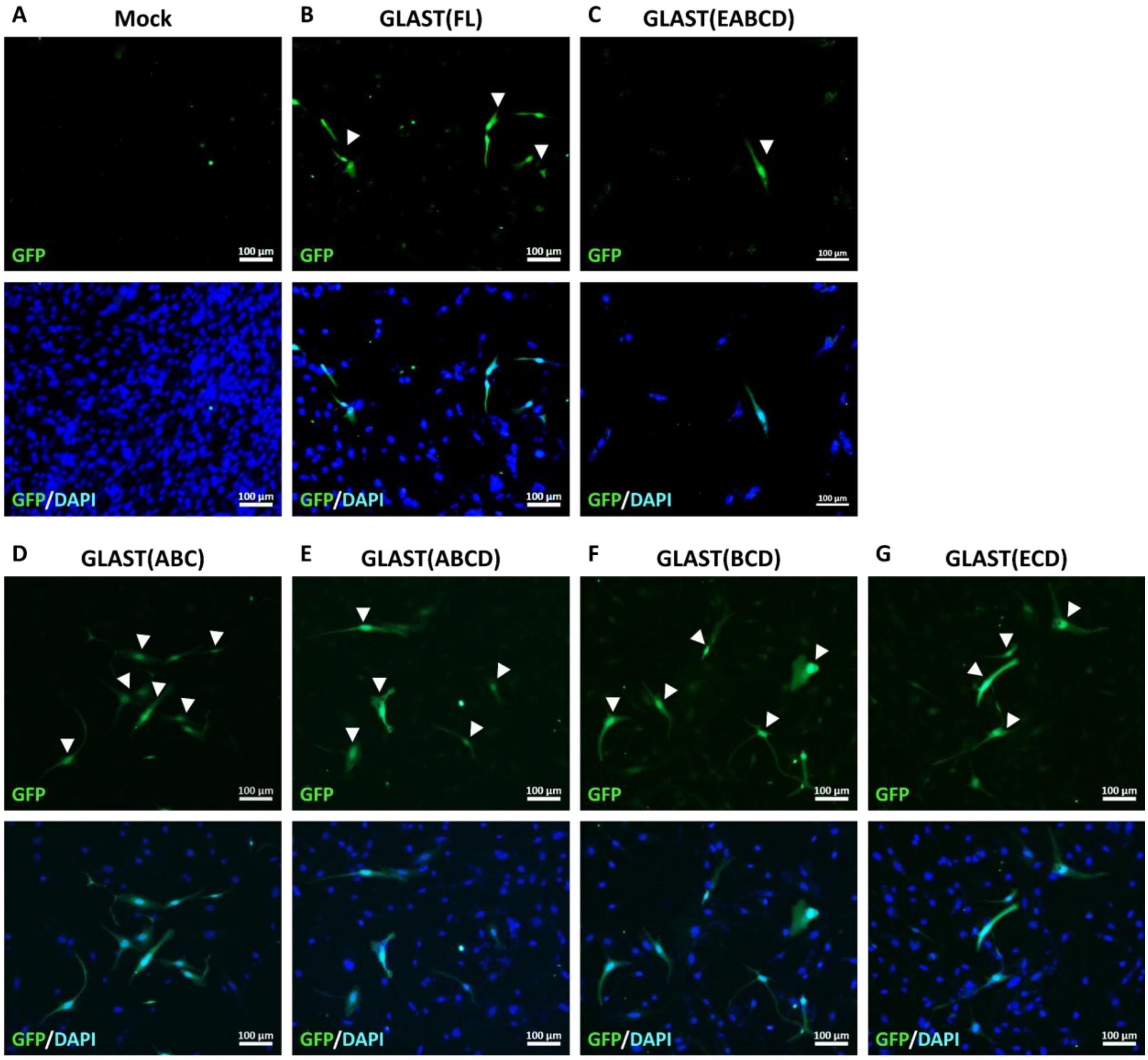


**Figure S3: *In vitro* testing of *GLAST* short promoters using AAV plasmids**. A-G) Transfection assay of AAV plasmids carrying the *GLAST* full-length (FL) and short promoters to test promoter activity *in vitro* using MIO-M1. White arrows indicate the GFP+ MIO-M1 cells. A) No transfection control (Mock), B) *GLAST*(FL) transfection and C-G) short *GLAST* promoters (EABCD, ABC, ABCD, BCD, ECD). Scale bar = 100 µm.

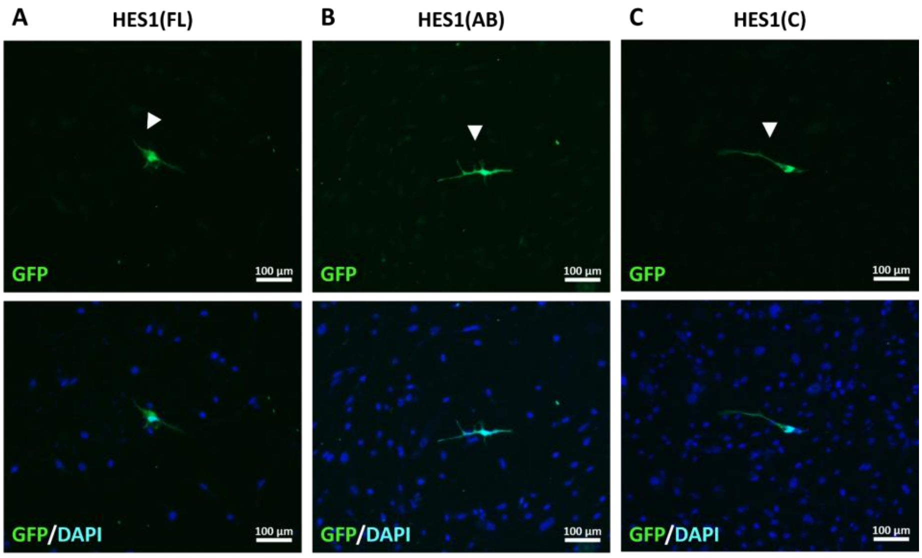


**Figure S4: *In vitro* testing of the *HES1* promoters using AAV plasmids.** Transfection assay of the AAV plasmids with A) *HES1* full-length promoter (FL), short B) *HES1*(AB) and C) *HES1*(C) promoters in MIO-M1 cells to assess promoter activity *in vitro*. Scale bar = 100 µm.

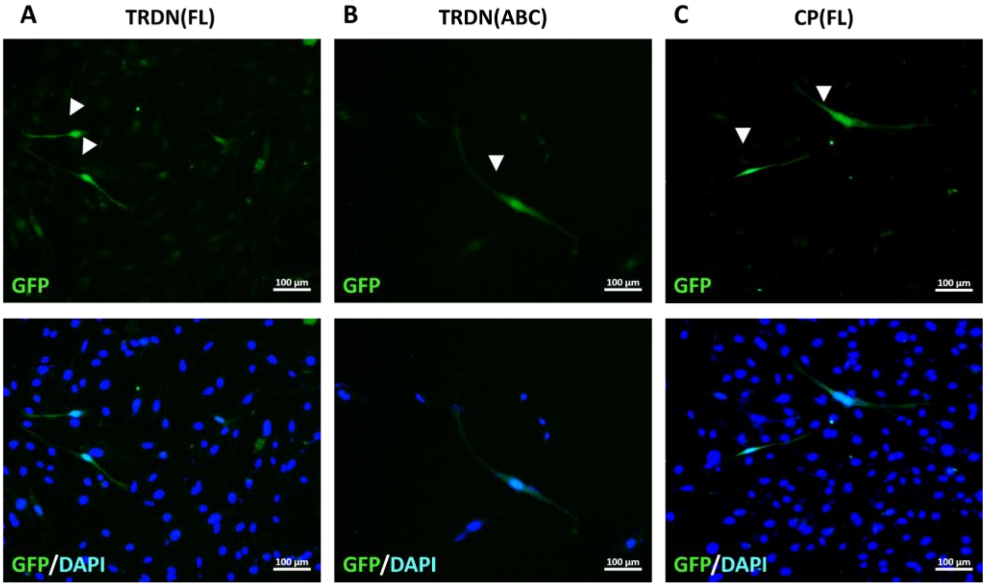


**Figure S5** *In vitro* testing of *TRDN* and *CP* human promoters. Transfection of the AAV plasmids with A) *TRDN* fulllength promoter (FL), B) *TRDN*(ABC) and C) *CP* promoter into MIO-M1 cells to assess promoter activity *in vitro*. Scale bar = 100 µm.
